# Integrated Genomic and Methylome Profiling Reveals Promoter Repression and Age-Linked CpGs in the California Mussel

**DOI:** 10.1101/2025.08.27.672726

**Authors:** Laurel S. Hiebert, Ava Soesbe, Tzung-Fu Hsieh, Qirui Cui, Soojin V. Yi

**Affiliations:** Department of Ecology, Evolution, and Marine Biology, University of California, Santa Barbara, CA, USA; Department of Plant and Microbial Biology, North Carolina State University, Raleigh, NC 27695, USA; Plants for Human Health Institute, North Carolina State University, North Carolina Research Campus, Kannapolis, NC 28081, USA; Department of Ecology, Evolution, and Marine Biology, Department of Molecular, Cellular and Developmental Biology, Neuroscience Research Institute, University of California, Santa Barbara, CA, USA

**Keywords:** Methylome, DNA methylation, promoter methylation, gene body methylation, epigenetic clock, age-associated methylation, phenotypic plasticity, population genomics, marine invertebrate, bivalve mollusk

## Abstract

While most knowledge of animal DNA methylation comes from vertebrates, this epigenetic mark remains poorly understood in invertebrates, which comprise the majority of animal diversity. For instance, how promoter and gene body methylation contribute to gene regulation, and how methylation relates to aging, are still relatively unknown in most invertebrates. Focusing on the California mussel (*Mytilus californianus*), we paired whole-genome resequencing and whole-genome bisulfite sequencing from the same individuals and evaluated relationships among promoter methylation, gene body methylation, gene expression, and age. Using seven individuals spanning a range of body sizes from the Santa Barbara Channel, California, we found standing genetic variation levels similar to related species and a relatively small effective population size. CpG methylation was enriched in gene bodies, and gene body methylation was positively associated with expression. Promoter methylation was less frequent but showed a strong negative association with expression and remained the best predictor of repression after accounting for gene body methylation, aligning with patterns widely documented in vertebrates and adding to the limited evidence in invertebrates that promoter methylation can be regulatory. We identified thousands of age-associated differentially methylated loci with directional changes across age classes, providing candidate sites for epigenetic clocks that could enable assessment of biological age, health, and stress resilience in wild and cultured populations.

## Introduction

The California mussel (*Mytilus californianus*) is an abundant species in rocky intertidal and subtidal ecosystems along the Pacific coast of North America, ranging from the Aleutian Islands to central Mexico (Sagarin and Somero, 2006; Soot-Ryen, 1955). As an “ecosystem engineer,” it stabilizes substrates, supports diverse intertidal communities (Gutiérrez et al., 2003), and has been an important food source for coastal human populations for millennia (Jones and Richman, 1995). Remarkably, *M. californianus* can live for several decades, possibly exceeding 100 years in low disturbance habitats (Suchanek, 1981). Across meters of tidal height and thousands of kilometers of coastline, it encounters steep local gradients and broad geographic variation in abiotic and biotic conditions. This includes aerial exposure during low tide and space competition for attachment surfaces at the shore scale, and regional differences in temperature regimes and shifts in predator communities across its range. These nested gradients and recurrent disturbances test the limits of both local adaptation and phenotypic plasticity.

*Mytilus* species have long served as models in ecological, physiological, and population genetic research. *M. californianus* was central to the formulation of the Keystone Species Concept (Paine, 1966), and, unlike some of its congeners, shows no known hybridization (Fraïsse et al., 2016), making it well-suited for genomic studies unconfounded by introgression. *M. californianus* exhibits year-round spawning (Young, 1946) resulting in release of millions of gametes per spawning event (White, 1937), high dispersal via a long-lived planktonic larval phase (Strathmann, 1987; Trevelyan and Chang, 1983), and broad environmental tolerance (Logan et al., 2012; Sagarin and Somero, 2006). These traits contribute to a pattern of panmixia and absence of significant population structure (Addison et al., 2008; Engel, 2004; Levinton and Suchanek, 1978; Ort and Pogson, 2007), which is a relatively rare phenomenon as most species exhibit at least some degree of population differentiation, even in species with long planktonic larval stages (Grosberg and Cunningham, 2001; Hellberg et al., 2002). Such genetic homogeneity over its more than 6000 km range has been hypothesized to constrain local adaptation and favor phenotypic plasticity (Levins, 1986; Sanford and Kelly, 2011).

*M. californianus* exhibits a range of phenotypically plastic traits, such as differences in shell morphology and calcification rate (Richards et al., 2025). Epigenetic regulation, particularly DNA methylation, is a plausible mechanism enabling such plasticity (Bogan and Yi, 2024; Roberts and Gavery, 2012) and may also play a role in age-related physiological changes. Such epigenetically mediated plasticity is especially important for sessile species that cannot behaviorally avoid environmental stressors. In *M. californianus,* prior work on circadian rhythms in gill tissue revealed extensive tidal-cycle–driven shifts in gene expression and metabolism, with methylation-machinery-linked protein abundance changes suggestive of a role for DNA methylation in mediating these responses (Connor and Gracey, 2011; Elowe and Tomanek, 2021). Combined with similar findings in other intertidal mollusks (Clark et al., 2018), this points to DNA methylation as a potential mediator of environmentally responsive plasticity in this species.

In invertebrates, DNA methylation is typically concentrated in gene bodies, a mosaic pattern distinct from the comparatively genome-wide methylation of vertebrates (Feng et al., 2010; Suzuki and Bird, 2008; Zemach et al., 2010). Gene body methylation (GbM) has been proposed to stabilize gene expression (Gatzmann et al., 2018; Huh et al., 2013; Li et al., 2018), regulate alternative splicing (Flores et al., 2012; Foret et al., 2012), positively influence expression (Gavery and Roberts, 2013), associate with constitutively expressed or conserved genes (Sarda et al., 2012; Wang et al., 2013), and facilitate environmental acclimation (Liew et al., 2018). However, several studies report no association between GbM changes and gene expression or expression plasticity (Abbott et al., 2024; Dixon and Matz, 2022).

Because in invertebrates DNA methylation is often concentrated in gene bodies, promoter methylation in invertebrates has historically received less attention. Recent evidence, however, shows promoter methylation can be widespread and functionally relevant in some invertebrate taxa (Keller et al., 2016). In bivalve mollusks, some studies have linked promoter methylation with transcriptional repression, similar to many vertebrates (Rivière, 2014; Saint-Carlier and Riviere, 2015). However, reported patterns are inconsistent: some find positive correlations between promoter methylation and expression (Olson and Roberts, 2014), while others find a mix of positive and negative associations (Fu et al., 2021; Tan et al., 2022).

When considering the potential adaptive significance of DNA methylation in mollusks, some studies detect no significant relationship between environmentally induced methylation changes and gene expression (Downey-Wall et al., 2020; Johnson et al., 2022; Rajan et al., 2021), while others report clear regulatory effects, including environmentally responsive regulation underlying phenotypic plasticity (Dang et al., 2023; Lim et al., 2021; Olson and Roberts, 2014). This mixed evidence highlights the need to examine promoter methylation and GbM together to clarify their relative and combined roles in regulating gene expression and environmental responsiveness.

Building on the available *M. californianus* reference genome (Paggeot et al., 2022), we pair whole-genome resequencing (WGS) and whole-genome bisulfite sequencing (WGBS) from adductor muscle of seven *M. californianus* collected subtidally in the Santa Barbara Channel, California. The adductor muscle varies in cross-sectional area with tidal height and shows documented physiological plasticity (Dowd and Jimenez, 2019), making it a suitable tissue for probing methylation linked to plastic responses. WGS enables us to mask CpG sites overlapping polymorphic SNPs and to estimate genome-wide genetic diversity. Using these SNP-aware methylomes, we characterize global and regional methylation patterns with attention to promoters and gene bodies, and relate methylation to gene expression using publicly available adductor muscle RNA-seq. We also leverage the relationship between ontogenic age and shell length in California mussels and validate length as an age proxy via calibration to shell growth-ring counts to ask whether age-associated epigenetic changes are detectable in this species. Because age commonly covaries with methylation in animals, often at specific CpGs that change predictably with age and support epigenetic “clock” inferences in vertebrates and several invertebrates (Brink et al., 2024; Fairfield et al., 2021; Hearn et al., 2021), we use shell length as a proxy for age to test age as a covariate of methylation variation and to identify candidate age-associated loci. By integrating population genomics with methylation profiling, we provide insight into the regulatory architecture of the *M. californianus* methylome and its potential contributions to plasticity, environmental responsiveness, and aging.

## Results

### A survey of nucleotide diversity in California mussels

To evaluate levels of standing genetic variation in *Mytilus californianus*, we performed whole-genome resequencing on seven individuals collected subtidally near Santa Barbara, California (for sequence data details, see Table S1). Across the ∼1.7 Gb genome (Paggeot et al., 2022), we identified 27,829,423 high-confidence single-nucleotide polymorphisms (SNPs), representing 1.63% of genomic sites.

Genome-wide nucleotide diversity (π) was estimated with two independent variant-calling pipelines: π = 3.29×10^−3^ from the BCFtools (Danecek et al., 2021) call set and π = 3.83×10^−3^ from the FreeBayes (Garrison and Marth, 2012) call set, yielding a mean 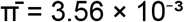. These values are comparable to estimates from the related mussel *M. coruscus* (3.21–3.79 × 10^−3^; (Guo et al., 2023)).

Using the molluscan germline mutation rate measured for white abalone (*Haliotis sorenseni*; μ = 8.60 × 10^−9^ mutations·bp^−1^·generation^−1^; (Wooldridge et al., 2025)), we estimated effective population size as 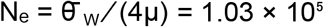 (95% range 0.80–1.46 × 10^5^ using μ’s reported CI). Caller-specific estimates were 9.56 × 10^4^ (BCFtools) and 1.11 × 10^5^ (FreeBayes).

To compare the effective population size with the census population size, we estimated the total number of *M. californianus* along the mainland Santa Barbara coast (Point Conception to Rincon) and the northern Channel Islands by multiplying rocky-intertidal area, regional mussel percent cover, and within-bed density.

Rocky-intertidal area, derived from USGS coastal baselines with representative bench widths, was set to 2.66, 6.31, and 11.05 km^2^ for low, mid, and high scenarios (Johnson et al., 2013; O’Neill et al., 2018). Percent cover was summarized from MARINe long-term photoplot surveys for sites in the region and bracketed at 0.10, 0.35, and 0.60 (Lohse et al., 2023; Richards and Rich, 2009). Within-bed density, drawn from regional measurements that typically range 10^2^–10^3^ m^−2^, was bracketed at 500, 1,500, and 3,000 m^−2^ (Blanchette and Gaines, 2007; Seed and Suchanek, 1992). The resulting intertidal census estimates were 1.3 × 10^8^ (low), 3.3 × 10^9^ (baseline), and 2.0 × 10^10^ (high) individuals. These values refer to the intertidal band only; the lower boundary of the mussel bed is set near the low-tide line by *Pisaster* predation. Subtidal populations were excluded from this estimate (Love, 2019; Paine, 1976). Therefore, the effective population size of California mussels is several orders of magnitude lower than the census population size, consistent with patterns across many taxa with high fecundity and variable reproductive success (Árnason et al., 2023; Hedgecock and Pudovkin, 2011).

### Genomic DNA methylation landscape of California mussels: global and regional methylation

We used whole-genome bisulfite sequencing (WGBS) to profile genomic DNA methylation in *M. californianus* (sequencing details in Table S1). To improve accuracy, we constructed SNP-aware methylomes by masking CpGs that overlapped segregating C↔T (reference strand) or G↔A (complement) variants identified from matched whole-genome resequencing. This filtering suppresses false hypomethylation at polymorphic sites and is recommended by methodological work and demonstrated in animal systems (Laine et al., 2023; Liu et al., 2012; Wu et al., 2020).

Across the seven individuals, the genome-wide methylation landscape was characteristic of mosaic invertebrate methylomes (Table S1). Mean %mCG across all CpGs across all seven samples was 11.5% (per-sample genome-wide averages in Table S2). We quantified methylation across gene bodies (TSS to TTS, including exons and introns) and in a proximal-promoter window defined as the 150 bp immediately upstream of the TSS, which is known to directly influence transcription initiation in metazoans (e.g., Carninci et al., 2006).

Both gene body methylation (GbM) and proximal promoter methylation were negatively correlated with CpG observed/expected ratios (CpG o/e) (Fig. 1A–B). GbM showed a strong negative correlation (Spearman’s ρ = –0.617, *P* < 10^-15^), while proximal promoter methylation showed a weaker but still negative relationship (ρ = –0.367, *P* < 10^-15^; Fig. 1A–B). These patterns are consistent with long-term maintenance of DNA methylation and evolutionary depletion of mutable CpGs (Bird and Taggart, 1980; Yi and Goodisman, 2009).

**Figure 1.**
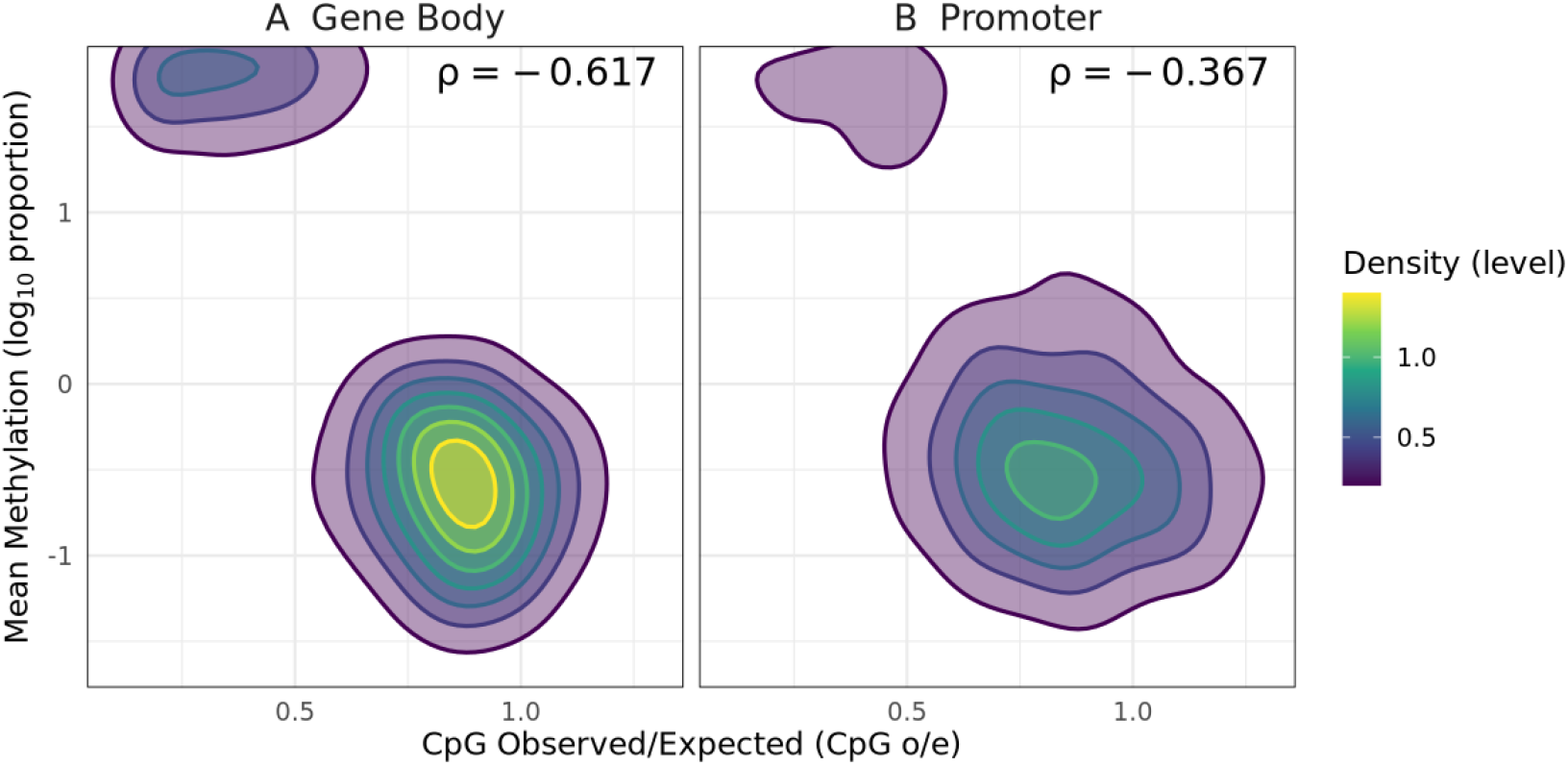
Relationship between CpG depletion and DNA methylation in gene bodies and promoters in *Mytilus californianus* genome. Bivariate density plots showing relationship between CpG observed/expected (CpG o/e) and mean DNA methylation (log_10_-transformed proportion) for **(A)** Gene bodies and **(B)** promoters of *Mytilus californianus*. Color intensity reflects smoothed two-dimensional density estimates with a square root transformation, and contour lines indicate isodensity levels to highlight cluster structure. Spearman’s rank correlation coefficients (ρ) are shown in the upper right of each panel, quantifying the strength and direction of association. Density and methylation values are computed per genomic element, including only regions with sufficient CpG content and coverage. Promoters are defined as spanning from the transcription start site (TSS) to –150 bp and are limited to coding genes. Gene body: ρ = –0.617, *p* < 2 × 10^−16^; Promoter: ρ = –0.367, *p* < 2 × 10^−16^.

We related methylation to transcription using adductor-muscle RNA-seq from *M. californianus* collected at a site several hundred km to the north of our samples (NCBI SRA PRJNA1011953 / SRP509094; SRX24640938; SRR29117084). This dataset is part of the California Conservation Genomics Project (Shaffer et al., 2022). For each expression group, we generated gene-aligned methylation curves (“metagene plots”) by rescaling gene bodies to a uniform number of bins, binning flanking regions at fixed distances, orienting genes by transcriptional strand, and averaging CpG methylation per bin across genes. These curves show higher methylation across gene bodies and lower methylation in proximal promoters for highly expressed genes (Fig. 2A), with the lowest proximal-promoter methylation in the highest expression stratum (Fig. 2B).

**Figure 2.**
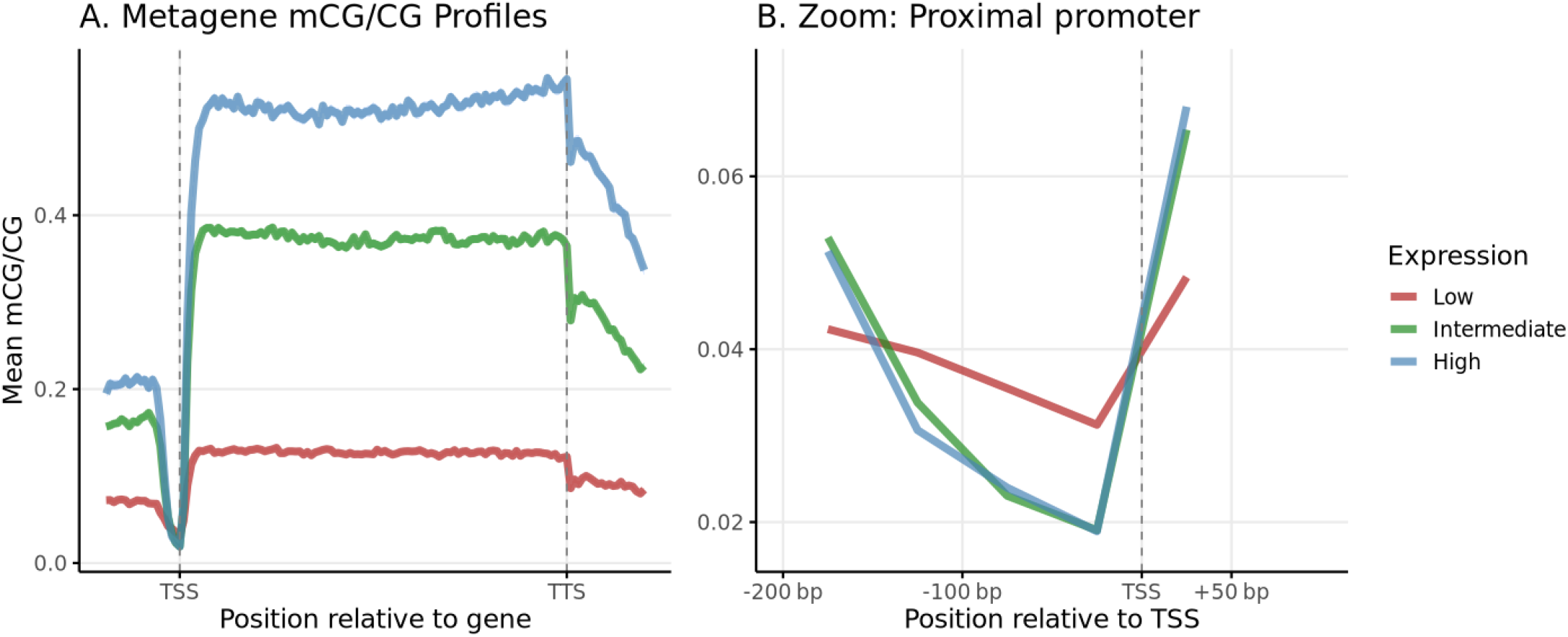
Relationship between gene expression level and DNA methylation across metagene and promoter regions of *Mytilus californianus*. **(A)** Metagene methylation profiles (mean mCG/CG) for genes grouped by expression level (Low, Intermediate, High), showing elevated gene body methylation in more highly expressed genes. Profiles span from upstream of the transcription start site (TSS) through to the transcription termination site (TTS), with methylation averaged across 100 equally sized gene body bins and 20 bins upstream/downstream. **(B)** Zoomed view of the proximal promoter region, showing reduced methylation immediately upstream of the TSS, especially in highly expressed genes. Lines represent smoothed averages across genes within each expression tertile. Only coding genes with sufficient methylation coverage and Salmon-derived expression estimates were included.

### Proximal promoter and gene body methylation interact to shape gene expression, with proximal promoter methylation having the dominant negative role

Even though most genes lacked promoter methylation, a substantial number harbored methylated proximal promoters. To evaluate its impact on transcription, we classified genes into four regional methylation states—high promoter/high body, high promoter/low body, low promoter/high body, and low promoter/low body—following the scheme of Keller et al. (2016) (Fig. 3A). In this classification, promoter methylation was computed in the proximal −150 bp window as we mentioned above. In our data this window size also maximized separation among expression groups, indicating that these regions are likely to impact transcription. Most genes fell into the low-promoter classes. The majority of genes (N = 11,842) exhibited little or no DNA methylation in either promoters or gene bodies and were generally lowly expressed (Fig. 3B). A large subset (N = 8,862) harbored gene-body methylation; within that subset, 689 also had high promoter methylation and 8,173 did not. Expression was highest for genes with low promoter and high gene-body methylation and lowest for genes with high promoter methylation regardless of gene-body state (Fig. 4B). Gene-scaled, TSS/TTS-anchored averages of CpG methylation, “metagene plots”, showed class-specific spatial patterns that paralleled these expression differences (Fig. 3C). Qualitatively similar rank-order relationships were observed if the promoter was instead defined as 2 kb upstream, though effect sizes were attenuated when averaging over the broader window.

**Figure 3.**
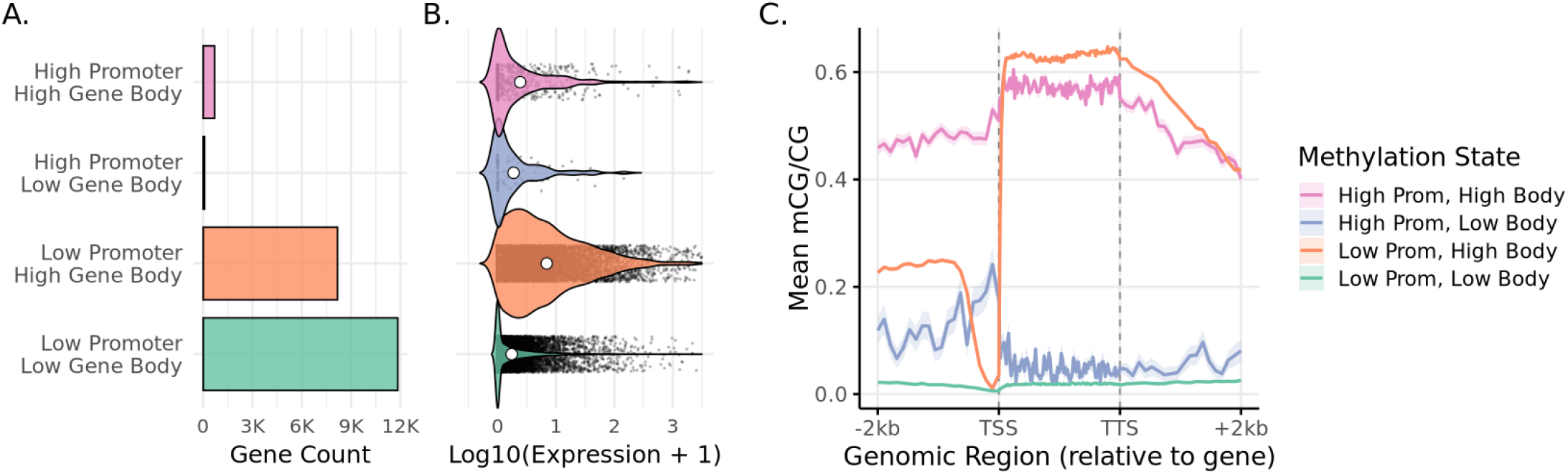
Combinatorial promoter and gene body methylation states and their relationship with gene expression and methylation profiles in *Mytilus californianus*. Genes were classified into four methylation states based on average DNA methylation levels in the proximal promoter (150 bp upstream of the transcription start site) and gene body. **(A)** The number of genes associated with each methylation state is shown, with bar colors matching those used throughout the figure. **(B)** Violin plots display the distribution of gene expression levels (log_10_[TPM + 1]) for each methylation state. Individual genes are shown as jittered points, and white circles represent group means. **(C)** Metagene plots show the average DNA methylation (mCG/CG) profiles across each methylation state, including 2 kb upstream of the TSS, the full gene body (scaled to a uniform length), and 2 kb downstream of the TTS. Dashed vertical lines mark the TSS and TTS.

**Figure 4.**
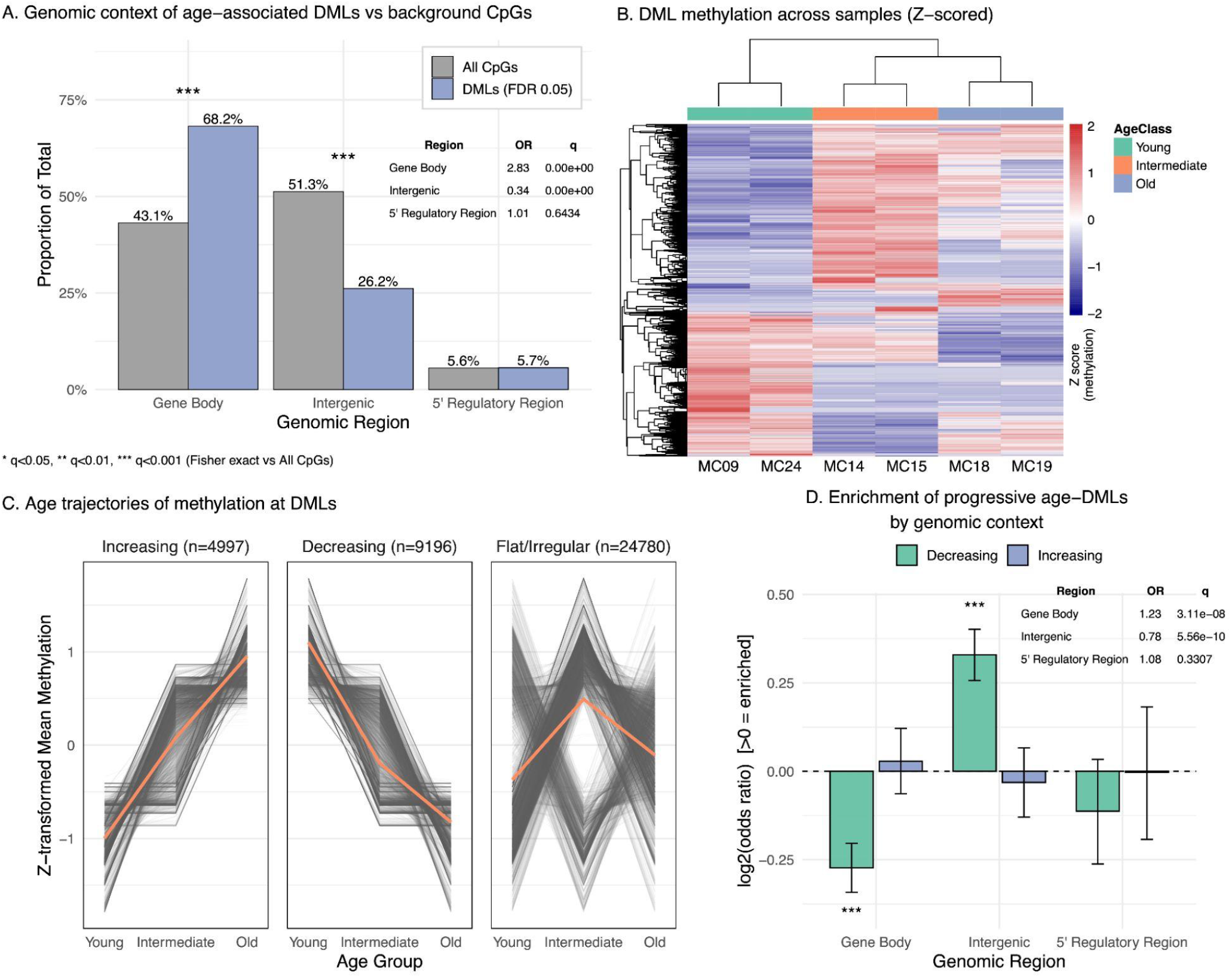
Genomic distribution and age-associated dynamics of DNA methylation at differentially methylated loci (DMLs). **(A)** Proportions of background CpGs (gray) and DMLs (purple; FDR ≤ 0.05) in the 5’ regulatory region, gene body, and intergenic regions. DMLs are enriched in gene bodies and depleted in intergenic regions relative to background. Background CpGs are covered sites from the BS-seq data under the same filters used to call DMLs. Enrichment was tested per region with Fisher’s exact test; the inset lists odds ratios (OR) and BH-adjusted q-values, and asterisks above bars denote significance (* q<0.05, ** q<0.01, *** q<0.001). **(B)** Heatmap of Z-transformed methylation at DMLs across individuals. Rows = loci; columns = samples with hierarchical clustering. Sample annotations indicate age group (Young, Intermediate, Old). The color scale (right) reflects Z-scores from low (blue) to high (red). **(C)** Age trajectories of methylation at DMLs across Young → Intermediate → Old, grouped by trend: Increasing, Decreasing, or Flat/Irregular. Gray lines show individual loci; thick orange lines are group means. Panel headers report the number of loci in each trend. **(D)** Enrichment of Increasing (purple) and Decreasing (teal) age-DMLs by genomic region, each compared to the full DML set at FDR ≤ 0.05. Bars show log2(odds ratio) with 95% CIs; values >0 indicate enrichment. Asterisks denote BH-adjusted significance as in (A).

To disentangle the independent contributions of the two regions, we analyzed only genes with measurable methylation in both places (mean mCG/CG ≥ 0.05 in promoter and gene body; N = 757). Partial Spearman correlations showed that proximal promoter methylation remained strongly and negatively associated with expression after adjusting for gene-body methylation (ρ = −0.178, p = 8.21×10^−7^), whereas the partial association for gene-body methylation (controlling for proximal promoter) was weak and not significant (ρ = 0.031, p = 0.391). Results were robust to reasonable inclusion thresholds for the both-methylated subset (for example, ≥ 0.001 and ≥ 1×10^−6^ in both regions; see Table S3). Together, these analyses indicate that while gene-body methylation is positively associated with expression, when present, proximal promoter methylation is the primary predictor of low expression and can override the positive association of gene-body methylation.

### Age calibration: shell length is a usable proxy for age

We used shell length as a proxy for ontogenetic age in *M. californianus*, an approach supported by classic growth studies and by a significant positive relationship between shell length and internal growth-band counts in this species (Coe and Fox, 1942; Denhel, 1956; Vriesman et al., 2021). Since growth rates are environmentally sensitive and vary by latitude, among other factors, we performed a preliminary calibration of the relationship between biological age and shell length in samples from the Santa Barbara Channel.

In a set of 19 mussels aged by growth-ring counts in the inner calcite layer of valve cross-sections (Table S4), shell length along the maximum growth axis showed a positive, monotonic relationship with age. A generalized additive model with a smooth on log(length) explained 89.1% of the deviance (adj. R^2^ = 0.873; edf = 2.56; F = 44.43; p < 2 × 10^−16^), with a residual RMSE of 0.83 years (Fig. S1). Predicted ages from this calibration were then used to assign the seven WGBS individuals to three classes: Young (MC09, MC24), Intermediate (MC11, MC14, MC15), and Old (MC18, MC19). Predicted ages for these samples ranged from 0.63 to 8.30 years (median 5.59 years), consistent with their observed shell lengths.

### DNA methylation trajectories change with age, and DMLs are concentrated in gene bodies

Using the Bioconductor package DSS(Feng and Wu, 2019; Feng et al., 2014; Park and Wu, 2016), we modeled methylation at each CpG as methylated vs unmethylated read counts in a beta–binomial generalized linear model with age class as a three-level factor. We tested the two adjacent contrasts (Young vs Intermediate and Intermediate vs Old) using Wald statistics and controlled the false discovery rate at 0.05 (Benjamini–Hochberg). CpGs significant in either contrast were called differentially methylated loci (DMLs). Loci significant in either contrast were combined to yield 38,973 differentially methylated loci (DMLs). DMLs were annotated relative to gene models and the 5’ regulatory regions (defined as 2000 bps upstream of TSS). Most DMLs mapped to gene bodies (68.2%), with fewer in intergenic (26.2%) and the 5’ regulatory region (5.7%) (Fig. 4A). Compared with the background distribution of covered CpGs, DMLs were enriched in gene bodies (Fisher’s exact test; OR = 2.83, BH-adjusted q < 0.05) and depleted in intergenic and 5’ regulatory regions (q < 0.05; Fig. 4A).

Z-scored methylation at DMLs largely separated samples by age class (Fig. 4B). Trajectory analysis across age groups showed 4,997 loci with increasing methylation, 9,196 with decreasing methylation, and 24,780 that were flat/irregular (Fig. 4C), indicating that age-related changes are widespread but potentially directional (i.e., ‘clock-like’) for a substantial subset of CpGs. Comparing genomic context between trend classes revealed a clear asymmetry: age-increasing DMLs are relatively gene-body–centric, whereas age-decreasing DMLs shift toward intergenic regions, with 5’ regulatory regions remaining a small and similar fraction across trends (Fisher’s tests vs all DMLs; Fig. 4D).

## Discussion

Whole-genome resequencing of seven *Mytilus californianus* individuals identified over 27 million SNPs, yielding genome-wide nucleotide diversity (π) estimates of 0.00383–0.00415 across two high-confidence call sets. These values fall within the intermediate range reported for marine bivalves, similar to *Mytilus coruscus* (0.00321–0.00379; (Guo et al., 2023)), and between published estimates for the Pacific oyster *Magallana gigas* (formally *Crassostrea gigas*), which vary widely by method (Li et al., 2021; Liu et al., 2025). Because π is sensitive to assembly quality, variant-calling pipelines, and filtering thresholds, cross-study comparisons should be interpreted cautiously.

Our estimate for effective population size (N_e_) of *M. californianus* is 1.19×10^5^–2.17×10^5^, far below the estimated census population size, which we bracketed between ∼10^8^ and ∼10^10^ individuals for the California coastline from Point Conception to Rincon, including the northern Channel Islands. This large N_e_-N gap is typical of many marine bivalves (reviewed by Plough, 2016). In *M. californianus,* previous work finds absence of geographic genetic structure, potentially due to high gene flow and/or weak spatially varying selection (Addison et al., 2008). Realized dispersal and recruitment, however, are shaped by larval behavior, coastal circulation, and early post-settlement mortality, so broad dispersal is relatively rare, even in broadcast spawners with long larval duration (Cowen and Sponaugle, 2009; Shanks, 2009). *M. californianus,* like many other bivalves, shows high heterozygote deficiency (Tracey et al., 1975; Zouros et al., 1988). These deficits have been interpreted as outcomes of high variance in reproductive success and sampling across temporally or spatially structured cohorts, that is, sweepstakes reproduction and Wahlund effects (Addison et al., 2008; Engel, 2004; Hedgecock, 1994; Tracey et al., 1975). At a site in California about 400km north of our sampling location, Engel estimated a contemporary N_e_ of 14,268 from allozyme data despite several hundred thousand to several million mussels present, paralleling our finding of low N_e_ relative to N (Engel, 2004). Together, these results suggest strong short-term variance in reproductive success that depresses contemporary N_e_ while long-term diversity remains high when averaged across generations.

Multiple phenotypic and life history traits vary geographically in *M. californianus*, including growth rate (Denhel, 1956; Smith et al., 2009), lifespan (Suchanek, 1981), and certain physiological traits (Logan et al., 2012). Such variation may reflect either local adaptation or phenotypic plasticity. Indeed, previous studies have reported plastic responses in *M. californianus* to environmental variation (Gleason et al., 2023), including developmental plasticity in thermal tolerance that depends on wave-exposure microhabitat (Gleason et al., 2018). Epigenetic processes, particularly DNA methylation, provide a plausible mechanistic basis for such plasticity (Bogan and Yi, 2024; Liew et al., 2018; Roberts and Gavery, 2012). Motivated by this, we examined how regional DNA methylation relates to gene expression and age in *M. californianus*.

Global CpG methylation in *M. californianus* averaged ∼11%, similar to recent reports from other mollusks (8.4–13.6% in bivalves; 9.5–16.6% in cephalopods; (Klughammer et al., 2023)). Gene body methylation (GbM) shows the canonical invertebrate pattern: enrichment within gene bodies and a positive association with expression (Mendoza et al., 2020; Zemach et al., 2010). Both promoter and gene body methylation were inversely related to CpG observed-to-expected ratios, consistent with long-standing methylation at these sites, although the strength of association differed between regions.

Against the limited and methodologically mixed evidence for invertebrate promoter methylation, *M. californianus* exhibits a relatively clear configuration: proximal promoter methylation is strongly and negatively associated with expression while GbM is an indicator of robust expression. This signal adds to evidence that promoter methylation can act as a transcriptional regulator in mollusks (Gavery and Roberts, 2013; Saint-Carlier and Riviere, 2015), even as mixed or context-dependent effects are reported in some invertebrates, including mollusks and tunicates (Fu et al., 2021; Tan et al., 2022). Clarifying whether the mussel pattern reflects lineage-specific elaboration or retention of an ancestral feature will require broader, phylogenetically informed sampling across protostome lineages.

Building on these baseline relationships between methylation and expression, we next examined age-related variation. Beyond baseline architecture, we identified over 38,000 differentially methylated loci associated with age, most within gene bodies, and a subset showed directional changes across three age classes. Similar age-associated methylation has been reported in other invertebrates, including lobsters and wasps (Brink et al., 2024; Fairfield et al., 2021), suggesting that even sparse invertebrate methylomes encode biologically meaningful age information. The identified loci provide a promising foundation for *M. californianus* epigenetic clocks, which, coupled with shell growth-ring validation and long-term field monitoring, could support assessment of biological age, healthspan, and environmental stress in wild or mariculture populations.

Viewed together, the baseline architecture and the age-linked changes point to complementary roles for the two methylation contexts. Proximal promoter methylation appears to set or lock expression state, consistent with promoter-linked repression, whereas GbM marks expressed genes and may provide a more dynamic substrate that modulates transcript output or splicing over the lifespan. The concentration of age-associated changes within gene bodies supports this interpretation and raises the hypothesis that GbM may provide primary candidates for epigenetic regulation of phenotypic plasticity and life-history variation, while promoter methylation imposes stronger constraints on whether genes are transcribed. These functional assignments remain hypotheses and will benefit from perturbation experiments and multi-tissue, longitudinal sampling.

Although this study was not designed to address sex-specific methylation, prior work indicates that sex and reproductive status can influence methylation landscapes in both vertebrates and invertebrates, including mollusks, even in somatic tissues (Teng et al., 2025). Given residual variability in our methylation data, incorporating sex as a covariate, sampling additional tissues, and increasing sample sizes will be important for disentangling the effects of sex and environment across the mussel’s lifespan.

### Conclusion

By integrating population genomics and methylome profiling in a marine invertebrate, this study reveals a methylation landscape with high regulatory potential and age-associated changes. The strong negative association between proximal promoter methylation and expression, coexisting with canonical GbM patterns points to diverse regulatory strategies. Age-associated methylation changes further suggest that the *M. californianus* methylome records aspects of individual life history that could be harnessed for aging and healthspan assessment. Together, these findings position *M. californianus* as a useful model for exploring the interplay among genetic diversity, epigenetic regulation, phenotypic plasticity, and aging in marine systems.

## Methods

### Sample Collection and DNA Extraction

Seven adult *Mytilus californianus* individuals were collected via SCUBA from the Santa Barbara Channel (California, USA). They were brought back and maintained in a flow-through seawater system. Adductor muscle tissue was dissected, flash-frozen in liquid nitrogen, and stored at –80 °C. Genomic DNA was extracted from ∼25 mg of tissue using a modified salting-out protocol: samples were incubated in lysis buffer (50 mM Tris-HCl pH 7.5, 400 mM NaCl, 20 mM EDTA, 0.5% SDS) with proteinase K and β-mercaptoethanol at 37 °C overnight. After mechanical disruption and precipitation of proteins and debris with 5 M NaCl and 3 M ammonium acetate, DNA was ethanol-precipitated, resuspended in TE buffer, treated with RNase A, and further purified using the Zymo DNA Clean & Concentrator-5 kit (Zymo Research, cat. no. D4004).

### Biological age determination and shell-length calibration

#### Growth-ring aging

We estimated biological age in a set of *Mytilus californianus* (n = 19) from annual growth bands in the inner calcite layer. Clean, oven-dried valves were covered with a layer of epoxy, and sectioned along the maximum growth axis (∼1.5 mm thick) with a tile saw. The sections were then each embedded in epoxy, mounted on glass slides, progressively sanded, and polished. Sections were examined under a dissecting microscope, and concentric bands were imaged and counted to obtain annual ring-based age (years), following classic and recent protocols (e.g., Coe and Fox, 1942; Denhel, 1956; Vriesman et al., 2021). For each individual, shell length along the maximum growth axis was measured to the nearest mm. The full calibration dataset (length and ring-based age) is provided in Table S4. Note that in many shells the band boundaries were indistinct and required interpretive judgment, and the protocol is still being refined. The calibration should therefore be regarded as provisional and used primarily for relative rather than absolute age inference, with no extrapolation beyond the observed size range.

#### Statistical calibration model

To relate age to size, we fit a generalized additive model (GAM) in R (package mgcv) with a smooth of the natural-log of length: *age (years) ∼ s(log(length(mm)), k = 4),* assuming Gaussian errors and an identity link, estimated by REML. The small basis dimension (k=4) was chosen a priori to avoid overfitting given n=19 while allowing mild curvature typical of decelerating shell growth. Model diagnostics (residual vs. fitted, QQ-plot, and concurvity) indicated adequate fit and no strong structure in residuals. Across the observed size range, the fitted relationship was positive and monotonic (Fig. S1).

#### Model performance

The GAM explained a large fraction of age variation (adj. R^2^=0.873; deviance explained = 89.1%). The smooth term had edf = 2.56 with F=44.43, p<2×10^-16^. The root-mean-squared error (RMSE) of age predictions was 0.83 years (Fig. S1).

#### Age prediction and class assignment for WGBS samples

We predicted ages for the seven whole-genome bisulfite sequencing (WGBS) individuals using the fitted GAM (predict.gam; point estimates shown as blue points in Fig. S1). Predicted ages ranged 0.63–8.30 years (median = 5.59 years). For downstream methylation analyses, samples were grouped into three age classes based on these predictions: Young (MC09, MC24), Intermediate (MC11, MC14, MC15), and Old (MC18, MC19). The calibration curve and 95% confidence band are shown in Fig. S1.

### Genomic Library Preparation and Short-read Whole-Genome Resequencing

Isolated DNA (1–2 µg) was sheared to ∼300 bp using a Covaris M220 sonicator (target BP = 300; peak incident power = 75 W; duty factor = 10%; cycles per burst = 200; treatment time = 90 s; 50 µL volume). Sheared DNA was cleaned with 1.2× AMPure XP beads (Beckman Coulter), end-repaired and A-tailed (NEBNext Ultra II DNA Library Prep Kit for Illumina, New England Biolabs), and ligated to Illumina-compatible multiplex adapters (NEBNext Multiplex Oligos for Illumina) following the manufacturer’s instructions. ∼5% of the adapter-ligated DNA was PCR-enriched with Q5 DNA polymerase (New England Biolabs) for 6 cycles. Enriched libraries were bead-cleaned and submitted for high-throughput sequencing at Psomagen (USA) on an NovaSeq X Plus (Illumina) platform to generate 150-bp paired-end reads.

Adapter trimming was performed using TrimGalore v0.6.10 (Krueger, Babraham Bioinformatics). Reads were aligned to the *M. californianus* reference genome (GCF_021869535.1; (Paggeot et al., 2022)) with Bismark v0.24.2 in non-bisulfite mode (Krueger and Andrews, 2011) using the Bowtie2 (Langmead and Salzberg, 2012) backend. BAM files were coordinate-sorted, deduplicated, and indexed using Samtools v1.9 (Li et al., 2009).

### SNP Calling

We used two complementary variant-calling strategies:

- *Approach 1* (fast, high-confidence): BCFtools v1.21 (Danecek et al., 2021) was used to generate genome-wide BCFs with mpileup (indel skipped, depth capped at 10,000), followed by SNP calling with bcftools call (multiallelic caller, indels excluded).
- *Approach 2* (haplotype-aware, high sensitivity): Reads were base-recalibrated with GATK v4 (DePristo et al., 2011; VanderAuwera and O’Connor, 2020), variants called per-sample with HaplotypeCaller, and joint genotyping done with GenotypeGVCFs. Filtering was applied on QD, FS, MQ, and SOR. In parallel, FreeBayes v1.3.2 (Garrison and Marth, 2012) was run across all samples and filtered for QUAL ≥ 20 and DP ≥ 10.

### Calculating diversity and effective population size (N_e_)

Genome-wide and windowed diversity metrics (π and Watterson’s θ) were computed using VCFtools v0.1.16 (Danecek et al., 2021). One sample with low WGS coverage was excluded in this analysis.

We calculated per-site Watterson’s θ (θ_W_) in non-overlapping windows and combined windows by callable-length weighting to obtain a genome-wide mean 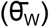. Genomewide diversity summary reported in Table S5. We converted diversity to effective population size assuming a diploid autosomal genome and neutrality, using 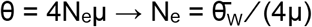. For μ we used the direct germline estimate from white abalone (*Haliotis sorenseni*), μ = 8.60×10^−9^ per site per generation (95% CI 6.10–11.11×10^−9^) (Wooldridge et al., 2025).

Where reported, confidence intervals on N_e_ reflect non-parametric block-bootstrap variance in 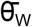 combined with mutation-rate uncertainty.

### Estimation of census size (N)

We estimated the number of *Mytilus californianus* in our region (from Point Conception

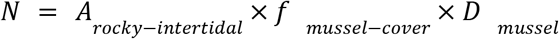

where *A_rocky-intertidal_* is the planform area of rocky intertidal habitat, *f_mussel-cover_* is the fraction of that area occupied by mussel bed, and *D_mussel_* is the density of mussels inside beds. We delineated *A_rocky-intertidal_* from USGS State Waters Map Series baselines and CoSMoS shoreline products, assigning representative bench widths by coastline segment to convert length to area (Johnson et al., 2013; O’Neill et al., 2018). We parameterized *f_mussel-cover_* from long-term MARINe rocky-intertidal percent-cover surveys using site time series for mainland and Channel Islands sites (Lohse et al., 2023; Richards and Rich, 2009). We drew *D_mussel_* from published measurements within *M. californianus* beds in the Point Conception region and syntheses of mussel-bed structure, which report typical densities on the order of 10^2^–10^3^ individuals m^−2^. We therefore used conservative brackets and did not convert cover to counts (Blanchette and Gaines, 2007; Seed and Suchanek, 1992). We constrained habitat to the intertidal zone where the mussel band occurs; the lower limit is set near low tide by *Pisaster* predation, although the species also occurs subtidally on hard substrates (Love, 2019; Paine, 1976).

### Whole-Genome Bisulfite Sequencing (WGBS)

WGBS libraries were constructed from the same seven individuals as the WGS data. Extracted gDNA was sheared with a Covaris M220 ultrasonicator, targeting ∼200 bp fragments, with fragment size confirmed via Agilent TapeStation. Libraries were prepared using the NEBNext Ultra II DNA Library Prep Kit (NEB, cat. no. E7645S) and NEBNext Multiplex Oligos (cat. no. 7535). Unmethylated lambda DNA (NEB N3013) was spiked in to assess conversion efficiency. Bisulfite conversion was performed with the EpiJET Bisulfite Conversion Kit (Thermo Scientific), followed by low-cycle PCR amplification with Q5U polymerase (NEB M0515S). Libraries were cleaned with AMPure XP beads (Beckman Coulter) and 150-bp paired-end reads were sequenced on an NovaSeqX (Illumina) platform.

### WGBS Read Processing and Global Methylation Estimation

Reads were adapter-trimmed with TrimGalore v0.6.10 (Krueger, Babraham Bioinformatics) and aligned using Bismark v0.24.2 (bisulfite mode) (Krueger and Andrews, 2011). Methylation extraction and deduplication were performed with Bismark’s deduplicate_bismark, bismark_methylation_extractor, and coverage2cytosine. Conversion rates (based on λ spike-in) exceeded 98.5% for all samples.

BSseq objects were constructed in R using the bsseq package (Hansen et al., 2012). SNP-masking was applied by removing loci overlapping high-confidence SNPs (from merged GATK-FreeBayes VCFs). Final global %mCG levels were computed per sample, and averaged across samples.

### CpG Observed/Expected (CpG o/e)

CpG o/e ratios were calculated for coding sequences (CDS) and proximal promoters (-150 bp to TSS) using custom Python scripts, following the formula: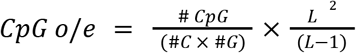 where L is the sequence length in nucleotides (Elango et al., 2009; Gavery and Roberts, 2010). Regions with no C or G were excluded. CpG methylation levels (from WGBS) were intersected with these features to calculate mean methylation, and correlation between CpG o/e and methylation was assessed using Spearman’s ρ.

### Gene Expression Quantification

To relate DNA methylation to transcription, we integrated WGBS with publicly available adductor-muscle RNA-seq from *M. californianus* collected at Moat Creek, Mendocino Co, CA (NCBI SRA BioProject PRJNA1011953 / Study SRP509094; Experiment SRX24640938; Run SRR29117084; PE100, 41.7 million read pairs; 8 Gb). Data was downloaded and quantified using Salmon v1.9.0 (Patro et al., 2017) in quasi-mapping mode. Gene-level TPMs were obtained via tximport (Soneson et al., 2015) using the RefSeq GFF annotation from the genome (GCF_021869535.1; (Paggeot et al., 2022)).

### Regional Methylation-Expression Analysis

For each gene, expression was computed as log_10_(TPM+1) and methylation. Mean mCG/CG was calculated for the proximal promoter (−150 bp to TSS) and for the gene body (TSS to TTS, exons plus introns) from one high-coverage WGBS sample (MC24). To focus on genes with measurable methylation in both regions, we restricted analyses to genes with promoter mCG/CG ≥ 0.05 and gene-body mCG/CG ≥ 0.05 (N = 757). We then estimated unique associations with expression using partial Spearman correlations.

- promoter vs expression | gene body: ρ = −0.178, p = 8.21×10^−7^
- gene body vs expression | promoter: ρ = 0.031, p = 0.391

Predictors were on their native 0–1 mCG/CG scale. P values are two-sided. Correlations were computed in R using the ppcor package (Kim, 2015; Kim and Yi, 2007). Results were robust to reasonable inclusion thresholds for the both-methylated subset; varying the minimum mCG/CG in each region from 0.05 to 0.001 or 1×10^−6^ yielded the same qualitative pattern (see Table S3).

### Methylation-state groups and length-scaled profiles

For descriptive summaries, genes were placed into four methylation-state groups using sample-wide medians for promoter and gene-body methylation: high-promoter/high-body, high-promoter/low-body, low-promoter/high-body, and low-promoter/low-body. For visualization, we generated length-scaled, gene-anchored averages of methylation across the promoter-to-terminator interval and flanking regions, aligned to TSS and TTS.

### Differential Methylation Analysis (DMLs)

Samples were grouped by age: Young (MC09, MC24), Intermediate (MC14, MC15), Old (MC18, MC19). MC11 was excluded due to lower WGBS coverage compared to the rest. DMLs were identified using DSS v2.43.2 (Feng et al., 2014; Park and Wu, 2016) in R with the DMLfit.multiFactor() and DMLtest.multiFactor() functions, modeling methylation as a function of AgeGroup. DMLs were filtered at p< 0.001 and FDR ≥ 0.1.

Genomic features were derived from the RefSeq GFF (GCF_021869535.1; (Paggeot et al., 2022)). A TxDb object was generated using GenomicFeatures and rtracklayer (Lawrence et al., 2013). Regions were defined as: proximal promoter (150 bp upstream of TSS), gene body (TSS to TTS, including exons and introns), intergenic (remaining loci). DMLs were annotated using GenomicRanges, and proportions per region were compared to background CpG distributions. Statistical enrichment of DMLs compared to background was assessed using Fisher’s exact test for each region.

We computed the mean and z-scored methylation for each retained CpG site per sample. CpGs were categorized by methylation trajectory across age classes using trend-based clustering into: increasing, decreasing, or flat/irregular patterns.

## Data Availability

Raw and processed WGS and WGBS data will be deposited in [repository] upon publication.

## Acknowledgments

We thank Cristoph Pierre (Director of Marine Operations at UCSB) for the mussel collection. This work was supported by a NSF grant (EF-2021635) and UCSB to SVY. We thank the members of the Yi lab, especially Ryan Son, Madison Weise and Haein Lee for discussion.

---

**Figure S1.**
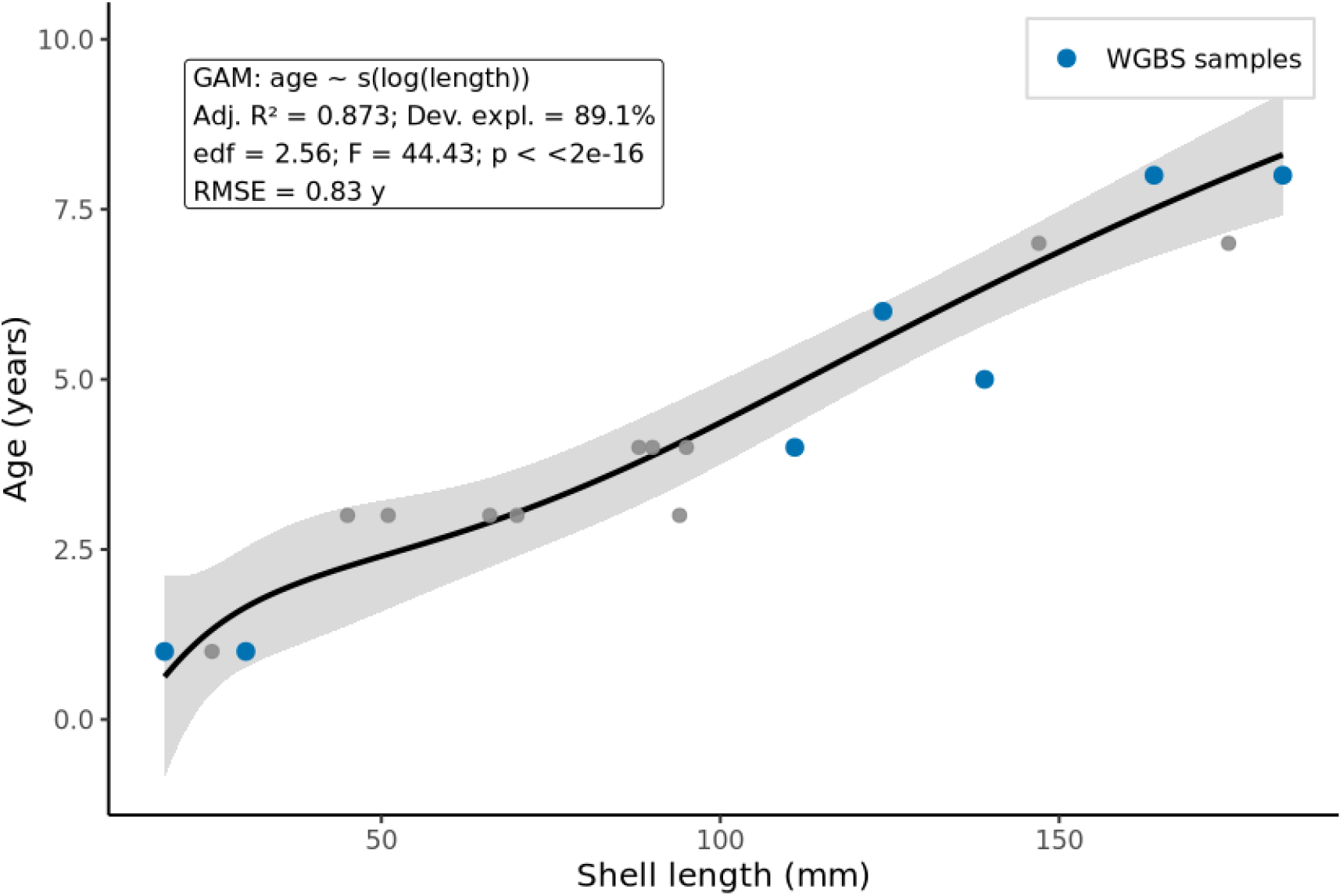
Calibration of age from shell valve length. A generalized additive model (GAM) relating age (years) to log shell length (mm) was fit to 19 mussels aged by growth-ring counts (gray points): age ∼ s(log(length)), k = 4. The black curve is the GAM fit and the gray band is the 95% CI; blue points mark the seven WGBS individuals. Model performance: adj. R^2^ = 0.873 (deviance explained = 89.1%), edf = 2.56, F = 44.43, p < 2 × 10^−16^; RMSE = 0.83 years. Predicted ages for WGBS samples (0.63–8.30 y; median = 5.59 y) were used to define age classes: Young (MC09, MC24), Intermediate (MC11, MC14, MC15), and Old (MC18, MC19). See Table S4 for calibration lengths and ring-based ages.

## Notes

### Competing Interest Statement

The authors have declared no competing interest.

