## Supplemental Tables for "Integrated Genomic and Methylome Profiling Reveals Promoter Repression and Age-Linked CpGs in the California Mussel"

| Table S1. Sequence data used in this study | | | | | | | |
|---|---|---|---|---|---|---|---|
| **Sequencing typ** | **Individual ID#** | **Tissue** | **Raw read pairs** | **Cleaned read pa** | **BS conversion r** | **Mapping efficie** | **Mean coverage (X)** |
| WGBS | 9 | Adductor muscle | 472,574,666 | 235,568,318 | 98.60% | 42.90% | 11.5 |
| WGBS | 11 | Adductor muscle | 488,680,790 | 243,409,955 | 98.70% | 12.40% | 3.6 |
| WGBS | 14 | Adductor muscle | 425,580,172 | 211,816,791 | 98.60% | 40.90% | 10 |
| WGBS | 15 | Adductor muscle | 276,816,596 | 138,314,727 | 99.70% | 45.20% | 9.2 |
| WGBS | 18 | Adductor muscle | 208,243,018 | 104,020,853 | 99.70% | 46.30% | 7 |
| WGBS | 19 | Adductor muscle | 202,222,662 | 100,996,626 | 99.70% | 45.70% | 6.8 |
| WGBS | 24 | Adductor muscle | 250,803,436 | 125,339,808 | 99.70% | 47% | 8.7 |
| WGS | 9 | Adductor muscle | 45,178,536 | 45,099,039 | NA | 11.90% | 0.85465 |
| WGS | 11 | Adductor muscle | 79,711,158 | 79,678,705 | NA | 46% | 5.59432 |
| WGS | 14 | Adductor muscle | 61,536,804 | 61,515,585 | NA | 45.90% | 4.21937 |
| WGS | 15 | Adductor muscle | 58,162,182 | 58,137,412 | NA | 46.10% | 4.14122 |
| WGS | 18 | Adductor muscle | 139,225,461 | 139,172,993 | NA | 46.10% | 9.66281 |
| WGS | 19 | Adductor muscle | 96,315,012 | 96,285,323 | NA | 46.70% | 6.85006 |
| WGS | 24 | Adductor muscle | 88,984,117 | 88,948,361 | NA | 45.90% | 6.11945 |

| Table S2. Average genome-wide %mCpG per sample | |
|---|---|
| **Sample** | **Genome-wide mGpG/CpG (%)** |
| M44-9-1yr | 12.522724 |
| M45-11-5yr | 13.865654 |
| M46-14-4yr | 12.736596 |
| M47-15-6yr | 11.343041 |
| M48-18-8yr | 10.312165 |
| M49-19-8yr | 10.023614 |
| M50-24-1yr | 9.972521 |

| |
|---|
| **Table S3. Sensitivity analyses for partial effects test** |
| Before filtering: total genes = 20,797. |
| Both regions ≥ 0.05: N = 757; partial ρ(promoter vs expression \| gene body) = −0.178, $p = 8.21\times10^{-7}$; partial ρ(gene body vs expression \| promoter) = 0.031, $p = 0.391$. |
| Both regions ≥ 0.001: N = 1,104; partial ρ(promoter \| body) = −0.269, $p = 9.19\times10^{-20}$; partial ρ(gene body \| promoter) = 0.213, $p = 8.48\times10^{-13}$. |
| Both regions ≥ $1\times10^{-6}$: N = 1,137; partial ρ(promoter \| body) = −0.258, $p = 8.81\times10^{-19}$; partial ρ(gene body \| promoter) = 0.226, $p = 1.3\times10^{-14}$. |

**S4. Preliminary age curve samples**

| Mussel Number | Length (mm) | Age (ring count) |
|---|---|---|
| 9 | 18 | 1 |
| 25 | 25 | 1 |
| 24 | 30 | 1 |
| 17 | 45 | 3 |
| 10 | 51 | 3 |
| 23 | 66 | 3 |
| 7 | 70 | 3 |
| 22 | 88 | 4 |
| 12 | 90 | 4 |
| 16 | 94 | 3 |
| 8 | 95 | 4 |
| 14 | 111 | 4 |
| 15 | 124 | 6 |
| 11 | 139 | 5 |
| 21 | 147 | 7 |
| 18 | 164 | 8 |
| 20 | 165 | 10 |
| 13 | 175 | 7 |
| 19 | 183 | 8 |

**Table S5. Genomewide Diversity Summary**

| PREFIX | AVG_PI | TAJIMA_D | SITES | THETA_RAW | THETA_NORM | GENOME_LEN |
|---|---|---|---|---|---|---|
| snps_filtered | 0.003287021241 | 0.2676 | 21389702 | 6726038.477 | 0.004071574673 | 1651950171 |
| freebayes_filt | 0.003830633703 | -nan | 9414696 | 9414696 | 0.005699140425 | 1651950171 |
